# RNA binding by Periphilin plays an essential role in initiating silencing by the HUSH complex

**DOI:** 10.1101/2024.07.17.602677

**Authors:** Stuart Bloor, Niek Wit, Paul J. Lehner

## Abstract

The human silencing hub (HUSH) complex is a transcription-dependent, epigenetic repressor complex that provides a genome-wide immunosurveillance system for the recognition and silencing of newly-integrated retroelements. The core HUSH complex of TASOR, MPP8 and Periphilin, represses these retroelements through SETDB1-mediated H3K9me3 deposition and MORC2-dependent chromatin compaction. HUSH-dependent silencing is RNA-mediated, yet no HUSH components contain any RNA-binding domain. Here we used an unbiased approach to identify which HUSH component was able to bind RNA and determine whether RNA-binding was essential for HUSH function. We identify Periphilin as the major RNA-binding component of the HUSH complex and show that Periphilin’s N-terminal domain is essential for both RNA binding and HUSH function. Periphilin binding to RNA was independent of its interaction with TASOR or MPP8, as its N-terminal domain was sufficient for RNA targeting. The artificial tethering of Periphilin to a HUSH-insensitive, nascent transcript, enabled the HUSH-dependent silencing of the transcript. This tethering of Periphilin allowed the RNA-binding region of Periphilin to be removed such that only its C-terminal domain was required for oligomerisation and interaction with TASOR. We therefore show that Periphilin is the predominant RNA-binding protein of the HUSH complex and this RNA-binding is essential for HUSH activity.

## Introduction

The success of endogenous retroviruses and retrotransposons in colonising the human genome is emphasised by the finding that together the retroelement family makes up more than 40% of the genome, while coding genes contribute less than 2% (1). Line-1s (L1s) alone constitute as much as 17% and are the only transcriptionally active retrotransposons in the human genome (2). Active retroelements encode their own endogenous reverse transcriptase which allows them to retrotranscribe their RNA to a cDNA copy which then integrates into the host genome. Whilst clearly vulnerable to retroelement invasion, genomes have also evolved potent mechanisms of chromatin defence which allow them to recognise and repress integrated nucleic acid and thus protect their genome from unwanted effects of integration (3).

Chromatin silencing pathways can be DNA- or RNA-directed. The DNA-binding KRAB-ZNF protein family constitutes the largest group of epigenetic transcriptional repressors with over 400 proteins, each of which binds their locus DNA in a sequence-specific manner. KRAB-ZNFs can then recruit TRIM28 (KAP1) which binds the SETDB1 histone methyltransferase and deposits H3K9me3 over the specific locus, resulting in the establishment of repressive heterochromatin (4). While the ZNF protein family provides an effective DNA sequence-directed silencing pathway it cannot recognise novel DNA sequences, i.e. newly integrated DNA which contains sequences to which it has not been previously exposed and are therefore not recognised by any ZNF family member. Silencing by the ZNF gene family is therefore learnt or acquired – it is not immediate or innate.

The ability to recognise and rapidly silence any newly integrated DNA species is particularly challenging as the repressor complex needs to recognise and silence new transgenes, whatever their sequence and wherever they integrate in the host genome, even when this occurs within the introns of actively transcribed host genes. The HUSH complex is an epigenetic transcriptional repressor complex whose essential function is to recognise and silence newly integrated retroelements, including both retroviruses and retrotransposons (5-8). We recently showed how HUSH uses the presence or absence of introns to distinguish, intron-containing ‘self-DNA’ from RNA-derived, and therefore by definition, intronless ‘foreign’ DNA (7).

Our current understanding of HUSH-mediated silencing suggests that the core HUSH complex of TASOR, MPP8 and Periphilin recognises and binds its target loci and recruits its effectors: the histone methyltransferase SETDB1 and MORC2, an ATP-dependent chromatin remodeller (8). SETDB1 deposits repressive H3K9me3, while the MORC2 ATPase compacts chromatin, which together leads to silencing of the specific locus. MPP8 encodes a chromodomain which binds H3K9me3 and methylated ATF7IP, the nuclear chaperone of SETDB1. However, neither the MPP8 chromodomain nor ATF7IP methylation are absolutely required for HUSH-dependent silencing (5,9) (10), implying other mechanisms must be required for both recruiting HUSH to chromatin and for sequence-specific silencing of its target loci.

The identified requirements for HUSH-dependent silencing are that newly integrated retroelements are: (i) transcriptionally active, (ii) at least 1 – 1.5 kB in length, (iii) adenine (A)- rich relative to transcription by RNA Pol II, and (iv) intronless (7). A requirement for active transcription in HUSH-dependent silencing originated from observations that HUSH preferentially silences evolutionarily young, full-length L1 elements, particularly young L1s located within transcriptionally permissive euchromatin (6,11). Subsequent experiments showed that decreased transcription, as a result of promoter deletion, dramatically reduces H3K9me3 deposition across the affected locus (7). Together, these observations strongly suggested that HUSH-dependent silencing requires active transcription and is therefore likely to be RNA-dependent. This finding was reinforced by the recognition that HUSH components share homology with the well-characterised RNA-induced transcriptional silencing (RITS) complex of S. pombe (9,12). In the case of RITS and other RNAi pathways, Argonaute-bound siRNAs direct RITS to chromatin via base-pairing interactions between siRNAs and nascent transcript (13,14).

For HUSH-dependent silencing to be RNA-directed, the initial recruitment of HUSH is likely to be dependent on the recognition of nascent RNAs and suggests that at least one component of the core HUSH complex should bind RNA, yet none of the HUSH proteins encode defined RNA-binding motifs. The likely RNA-binding candidate of the HUSH complex is Periphilin, a 55kD insoluble nuclear protein, which was identified as an mRNA-binding protein in an interactome screen in human cells (15-17). The C-terminus of Periphilin forms a dimer which binds TASOR and is required for HUSH complex assembly (18). Periphilin has a disordered, self-aggregating N-terminal domain (NTD) which is also essential for HUSH activity. This NTD is rich in arginine and tyrosine residues which likely accounts for its marked insolubility and has so far precluded further *in vitro* studies on RNA binding. From a functional viewpoint, this NTD can be partially complemented by low-complexity regions from other RNA-binding proteins (18). Our RNA immunoprecipitation sequencing (RIP-Seq) experiments supported these findings and provided genome-wide data that Periphilin binds transcripts from HUSH target loci and was therefore likely to play a critical role in HUSH chromatin localization, most likely through stabilising HUSH at its target loci (7).

However, RNA binding by both MPP8 and TASOR has also been reported (9,12,15,17-19), making it essential to establish which of the HUSH components can bind RNA, the nature of the bound RNA and the requirement for RNA binding in the recruitment of HUSH to chromatin and subsequent HUSH-dependent silencing. We therefore took an unbiased, systematic approach to identify which HUSH component(s) was RNA-binding and determine its role in recruitment of HUSH to chromatin and HUSH-mediated silencing.

We here show that (i) Periphilin is the main RNA binding component of the HUSH complex (ii) Periphilin’s N-terminal 127 amino acids are required for RNA binding and essential for HUSH function, and (iii) the RNA-binding requirement of full-length Periphilin can be bypassed by direct recruitment of Periphilin to nascent transcript such that only the minimal 110 amino acid CTD of Periphilin is required for HUSH-dependent silencing. This artificial recruitment of Periphilin to transcript enables the silencing of both a fluorescent reporter as well as an endogenous locus in a HUSH-dependent manner. Furthermore, RIP-seq analysis with Periphilin and Periphilin mutants shows that they bind HUSH RNA substrates even when not assembled in the HUSH complex.

## Materials and Methods

### Cell Culture

All Jurkat (ATCC), HEK293T (ATCC), HAP1 (Horizon) and HeLa (ECCAC) cell lines were maintained in Iscove’s Modified Dulbecco’s Medium (Gibco) supplemented with 10% FBS (Gibco) and 100U/ml penicillin, 100μg/ml streptomycin (Sigma). Cell cultures were routinely tested and found to be negative for mycoplasma infection (MycoAlert, Lonza).

### NanoLuc^®^ RNA Interactome Capture (NL-RIC)

This method was modified from the RNA interactome capture method developed by Castello et al. (16). Lentiviral transduced cells expressing candidate proteins fused to NanoLuc^®^ luciferase, were grown in duplicate plates to approximately 95% confluency, washed twice with PBS and one replicate of each plate was UV treated (254 nm, 300µJ/cm^2^) in PBS. Cells were harvested and cell pellets were lysed in a modified RIPA buffer (mRIPA) (1x TBS (25 mM Tris-HCl (pH 7.4), 130 mM NaCl, 2.7 mM KCL), 1% IGEPAL CA-630, 0.5% Sodium deoxycholate, 0.1% SDS, 1 mM EDTA, 1x cOmplete protease inhibitor cocktail (ROCHE), 20U/ml TURBO DNase (Invitrogen), 400 U/ml RNasin Plus (Promega)) and incubated 5 min at 37°C. Lysates were clarified by centrifugation and 2% of supernatant retained as input samples prior to the addition of oligo (dT) beads. Oligo (dT) pulldowns were incubated with rotation for 1h at 4°C, before washing the beads with; 1x mRIPA, 1x RIC lysis buffer (20 mM Tris-HCl (pH 7.5), 500 mM LiCl, 0.5% LiDS (wt/vol, stock 10%), 1 mM EDTA, 0.1% IGEPAL CA-630), 1x RIC buffer 1 (20 mM Tris-HCl (pH 7.5), 500 mM LiCl, 0.1% LiDS (wt/vol), 1 mM EDTA, 0.1% IGEPAL CA-630), 1 x RIC buffer 2 (20 mM Tris-HCl (pH 7.5), 500 mM LiCl, 1 mM EDTA, 0.1% IGEPAL CA-630), 1x RIC buffer 3 (Mix 20 mM Tris-HCl (pH 7.5), 200 mM LiCl, 1 mM EDTA) and eluted for 5 mins at 55°C in RIC elution buffer (20 mM Tris-HCl (pH 7.5) and 1 mM EDTA). Input samples diluted 1:50 in elution buffer, and eluates, were measured for NL luciferase levels in a plate reader using Nano-Glo® reagent (Promega). Fold enrichment was determined by dividing the percentage of input pulled down in the plus UV sample, by the percentage of input pulled down in the minus UV sample.

### Lentiviral production and transduction

HEK293T cells were transfected with a mixture of lentiviral vector, pCMVΔR8.91 and pMD2.G (5:4:1) using TransIT-293 transfection reagent (Mirus) at a 1µg DNA:2.25µl ratio, according to the manufacturer’s instructions. Lentiviral supernatants were harvested 48 h post transfection and either used fresh or stored at -80°C. Transduced cells were selected with the following drug concentrations: 2 μg/ml puromycin, 5 μg/ml blasticidin. 293T, HeLa cells 100 μg/ml hygromycin and Jurkat cells 1000 μg/ml hygromycin.

### Antibodies

Antibodies for immunoblotting: rabbit anti-TASOR (Atlas, HPA006735, 1:5,000), rabbit anti-MPP8 (Proteintech, 16796-1-AP, 1:10,000), rabbit anti-Periphilin1 (Sigma-Aldrich, HPA038902, 1:5,000), rabbit anti-SETDB1 (Proteintech, 11231-1-AP; 1:5,000), mouse anti FLAG (M2, Sigma-Aldrich, F3165, 1:10000), mouse anti-haemagglutinin (HA.11) tag (16B12, Covance, MMS-101P, 1:20,000), mouse anti-β-actin peroxidase conjugate (Sigma-Aldrich, A3854; 1:20,000). Secondary antibodies for immunoblotting: Horseradish peroxidase (HRP)-conjugated AffiniPure goat anti-mouse IgG (H+L) (Jackson ImmunoResearch, 115-035-146, 1:10,000), Antibodies for FACS analysis: mouse anti-MHC (W6/32 hybridoma S/N), Secondary antibody for FACS: Donkey anti-mouse IgG (H+L) Alexa Fluor™ 647 (Thermo, A32787). Antibodies for ChIP-qPCR: rabbit anti-H3K9me3 (Abcam, ab8898), rabbit anti-H3 (Abcam, ab1791).

### Western blotting

Cell pellets were lysed in 1x SDS sample buffer, plus Benzonase (Sigma) and incubated for 5 mins at 37°C. Lysed samples were heated for 10 mins at 90°C, prior to running on a NuPAGE 4-12% gel (Invitrogen). Gels were transferred onto PVDF membranes (Millipore), blocked in 5% milk in PBS, 0.2% Tween-20 and incubated overnight with primary antibody in blocking solution. Blots were imaged using the iBright CL1000 Imaging System (Invitrogen), and Supersignal ECL, West Pico Plus or West Dura reagents (Thermo Scientific).

### Flow cytometry

Live cells were analysed on an LSR Fortessa (BD). Data were analysed using FlowJo v10.10.0 software (BD).

### CRISPR–Cas9 mediated beta-2-microglobulin and ZNF37A gene knock-ins

For knock-in of a 0 or 12x BoxB-IRES-blasticidin (*bsd*) cassette into the 3’UTR of the beta-2-microglobulin gene (β2M) of Jurkat cells, they were transfected with plasmid pSpCas9(BB)- 2A-Puro (Addgene, #48139) containing an sgRNA targeting the sequence just downstream of the stop codon (exon 4) and either donor plasmid (a) 350 bp 5’ARM-(empty)-loxP-IRES-blasticidin-SV40 pA-loxP-350 bp 3’ARM or (b) 350 bp 5’ARM-12x BoxB loops-loxP-IRES-blasticidin-SV40 pA-loxP-350 bp 3’ARM. Transfected cells were enriched by blasticidin selection and single cell cloned. Clonal populations were validated by PCR on genomic DNA. sgRNA and donor sequences are listed in the Resource Table.

### UV-crosslinked RIP-seq

Cells were grown to 95% confluency, washed twice with PBS and UV treated (254 nm, 300µJ/cm^2^) in PBS. Cells were harvested and lysed in HLB-N buffer (10 mM Tris-HCl pH7.5, 10mM NaCl, 2.5 mM MgCl2 and 0.5% IGEPAL CA-630, 80 U/ml RNasin Plus, 1x cOmplete protease inhibitor cocktail), incubated on ice for 5 min and the lysate was then underlaid with 1/4 volume of HLB + NS (10 mM Tris-HCl pH7.5, 10mM NaCl, 2.5 mM MgCl2 and 0.5% IGEPAL CA-630, 10% (w/v) sucrose). Nuclei were pelleted by centrifugation (420xg, 5 min) and lysed in RIP buffer (25mM Tris pH 7.4, 150mM KCl, 5mM EDTA, 0.5% IGEPAL CA-630, 80 U/ml RNasin Plus). The nuclear fraction was sheared by sonication (Diagenode Pico), and treated with 4U TURBO-DNase, before the insoluble material was removed by centrifugation. The soluble fraction was immunoprecipitated with anti-HA magnetic beads (Pierce) for 2 h at 4°C. Beads were washed once with RIP buffer and the residual DNA was removed in by digestion with 2U TURBO DNase, in 1x TURBO DNase buffer, 5 min at 37°C. The beads were washed twice with RIPA buffer (50 mM Tris-HCl pH 7.4, 100 mM NaCl, 1% IGEPAL CA-630, 0.5% sodium deoxycholate, 0.1% SDS), once with high-salt RIPA (50 mM Tris-HCl pH 7.4, 500 mM NaCl, 1 mM EDTA, 1% IGEPAL CA-630, 0.5% sodium deoxycholate, 0.1% SDS) and once with low-salt wash (15 mM Tris-HCl pH 7.4, 5 mM EDTA), each wash included a 5 min incubation at room temperature with rotation. Beads were digested with proteinase K in proteinase K buffer (50 mM Tris-Cl pH 7.5, 100 mM NaCl, and 1 mM EDTA, 0.25% SDS) and RNA was isolated by standard phenol-chloroform extraction and GlycoBlue™ (Invitrogen) coprecipitation. Immunoprecipitated RNA was subjected to DNA library preparation using SMARTer Stranded Total RNA-Seq Kit V3—Pico Input Mammalian (Takara Bio) according to the manufacturer’s instructions with initial fragmentation at 94°C for 4 min and including the ribosomal RNA depletion step. The library quality was determined using Bioanalyzer, and sequenced on Illumina MiniSeq platform as paired-end 32-bp and 43-bp reads using MiniSeq High-Output 75 cycles kit.

Bioinformatics data processing and analyses were performed as follows: unique molecular identifiers (UMIs) were extracted with UMI-tools (10.1101/gr.209601.116). Reads were then trimmed with cutadapt (10.14806/ej.17.1.200) to remove adapter sequences. HISAT2 (10.1038/s41587-019-0201-4) was used to align the trimmed reads to the human genome (GRCh38). BigWig files were generated for each replicate using bamCoverage from deepTools (10.1093/nar/gkw257) with the CPM normalisation method. For visualisation: (i) mean bigWig files were generated from both replicates using wiggletools write (mean) (10.1093/bioinformatics/btt737) and wigToBigWig (10.1093/bioinformatics/btq351), and (ii) signal at various genomic locations was plotting using pygenometracks (10.1093/bioinformatics/btaa692). TE annotations (GTF format) were obtained from TEtranscripts (10.1093/bioinformatics/btv422).

Differential binding analysis was performed by: (i) selecting only mapped fragments less than 500 bp using a combination of Samtools view (10.1093/bioinformatics/btp352) and the Awk programming language (https://www.gnu.org/software/gawk/manual/gawk.html) (ii) sites of enrichment were identified in each replicate using the MACS2 (10.1186/gb-2008-9-9-r137) command callpeak with the settings --broad and --broad-cutoff 0.1 (iii) finally, the R package DiffBind (10.1038/nature10730) was used to determine the differential sites and create the PCA and box plots, using the alignment files from HISAT2 and the MACS2 peak files from each replicate.

TE analysis was performed as follows: (i) consensus peaks were obtained from both replicates using the bedtools (10.1093/bioinformatics/btq033) intersect command. The resulting BED file was then annotated to the nearest TSS to obtain gene names using the R packages GenomicFeatures (10.1371/journal.pcbi.1003118), ChIPseeker (10.1093/bioinformatics/btv145) and rtracklayer (10.1093/bioinformatics/btp328) (ii) genomic coordinates from these genes were extracted from the GRCh38 GTF file using a combination of grep (https://www.gnu.org/software/grep/) and awk and converted to the BED format with the BEDOPS gtf2bed command (10.1093/bioinformatics/bts277). BED files containing the locations of various classes of TE elements were generated with grep to subset the TE annotation GTF file described above, and converted to the BED format with BEDOPS gtf2bed. To count how many genes contained TE elements, BEDTools intersect (10.1093/bioinformatics/btq033) with the –c option was used using the BED file containing PPHLN1-bound genes coordinates (-a) and the BED files containing TE genomic coordinates (-b). Calculation of the percentages and plotting was performed with the R packages Tidyverse (10.21105/joss.01686) and Cowplot (https://github.com/wilkelab/cowplot). Tukey multiple comparisons of means were performed to test whether the genes bound by the Periphilin variants had different levels of TE content compared to all genes in the genome.

### Chromatin immunoprecipitation

10^7^ cells were cross-linked for 10 min in 1% formaldehyde, quenched for 5 min in 0.125 M glycine, washed in PBS and lysed in cell lysis buffer (1 mM HEPES, 85 mM KCl and 0.5% NP-40). Nuclei were pelleted by centrifugation and lysed in MNase buffer (10 mM Tris-HCl pH 7.4, 10mM NaCl, 3 mM MgCl2, 1mM CaCl2, 10 mM EDTA and 4% IGEPAL CA-630) for 10 min. Chromatin was digested with limited MNase treatment, stopped by the addition of Nuclei lysis buffer (50 mM Tris-HCl pH 8.0, 10 mM EDTA, 10mM EGTA, 1% SDS) and sheared in a Bioruptor (Diagenode Pico) to obtain a mean fragment size of approximately 300 bp. Insoluble material was removed by centrifugation and the samples supernatants were diluted to a final concentration of 0.1% SDS. Samples were precleared with Protein G magnetic beads (Pierce) and then immunoprecipitated overnight with 5 μg primary antibody and Protein G magnetic beads. Beads were washed twice with wash buffer 1 (20 mM Tris-HCl pH 8.0, 2 mM EDTA, 50 mM NaCl, 1% Triton X-100, 0.1% SDS), once with wash buffer 2 (10 mM Tris-HCl pH 8.0, 1 mM EDTA, 250 mM LiCl, 1% NP-40, 1% sodium deoxycholate) and twice with TE. Protein-DNA complexes were eluted in 100 mM NaHCO3 and 1% SDS at 65°C. The NaCl concentration was adjusted to 300mM, RNaseA was added, and cross-links reversed by overnight incubation at 65°C. The samples were then Proteinase K digested for 2 h at 45°C, before spin column purification (Qiagen PCR Purification Kit) of the DNA. Quantification by qPCR was performed on a QuantStudio 6 Flex Real-Time PCR System (Thermo Fisher Scientific) using SYBR green PCR mastermix (Thermo Fisher Scientific). qPCR primer sequences are listed in the Resource Table.

### Plasmid constructs

Non-intronic 3’UTR reporter plasmids containing 0, 6, 12 or 18 BoxB or MS2 stem loops, were generated from plasmid pHRSIN-P_SFFV_-iRFP^STOP^(BamHI-NotI)-WPRE-P_GK_-Hygro or pHRSIN-P_SFFV_-NanoLuc^STOP^(BamHI-NotI)-WPRE-P_GK_-Hygro, by sequential insertion of 6x BoxB or 6x MS2 loop sequences containing internal BamHI-NotI cloning sites. Internal intronic reporter plasmids LentiREV-(SV40pA-iRFPi(NxBoxB)-P_SFFV_)-P_SV40_-Bsd-WPRE were generated by the silent insertion of a consensus U2-dependent splice site (20) into either the NanoLuc or iRFP_713_ coding sequence, ligated into LentiREV-(SV40pA -GFP-P_SFFV_)-P_SV40_-Bsd-WPRE, replacing the GFP (BamHI-NotI). As with the 3’UTR constructs 0, 6, 12 and 18 stem loops, were generated by the sequential insertion of 6x BoxB or 6x MS2 loop sequence oligos. LentiREV-(SV40pA-iRFP_713_-P_SFFV_)-P_SV40_-Bsd-WPRE was generated by replacing the GFP in plasmid LentiREV-(SV40pA-GFP-P_SFFV_)-P_SV40_-Bsd-WPRE with an iRFP_713_ cDNA (BamHI-NotI). The iRFP_713_ 5’UTR and 3’UTR intronic constructs were generated from LentiREV-(SV40pA-iRFP_713_-P_SFFV_)-P_SV40_-Bsd-WPRE by insertion of a PCR product containing the 12x BoxB fragment of into either the BamHI site for 5’UTR or SalI site for 3’UTR.

## Results

### Periphilin is the primary RNA binding protein of the HUSH repressor complex

To characterise which HUSH complex protein(s) might bind RNA in the HUSH complex we took an unbiased, systematic approach. Initially, we used a modified version of the RNA Interactome Capture assay (RIC) (16), NanoLuc^®^ RNA Interactome Capture (NL-RIC), to compare binding of each of the three core HUSH components to cellular mRNAs. Like conventional RIC, NL-RIC uses UV crosslinking and oligo(dT) mediated RNA pulldown to isolate polyadenylated mRNA and its associated crosslinked RNA binding proteins (RBPs). We complemented HUSH component knockout HEK293T cells with their respective N-/C-terminal NanoLuc^®^ luciferase (NL) tagged HUSH components and performed NL-RIC pulldowns. Following normalisation against input levels, we divided the eluate NL levels of the UV treated samples by those in their respective UV untreated samples to obtain a fold enrichment in NL binding, which is used as the proxy for mRNA binding **(Fig. 1A**). The empty N-terminal NL vector was used as the negative control and NL N-terminal tagged hnRNP-C (NL:hnRNP-C) as the positive control. Both N- and C-tagged Periphilin were the only HUSH components enriched on transcript pulldowns (**Fig. 1B**), with neither N-/C-tagged TASOR nor MPP8 showing any significant enrichment over the control samples (**Fig. 1B**). With the caveat that our NL-RIC assay utilises processed mRNA for the pulldown, these data suggest that Periphilin is the dominant RBP in the HUSH complex.

**Fig. 1.**
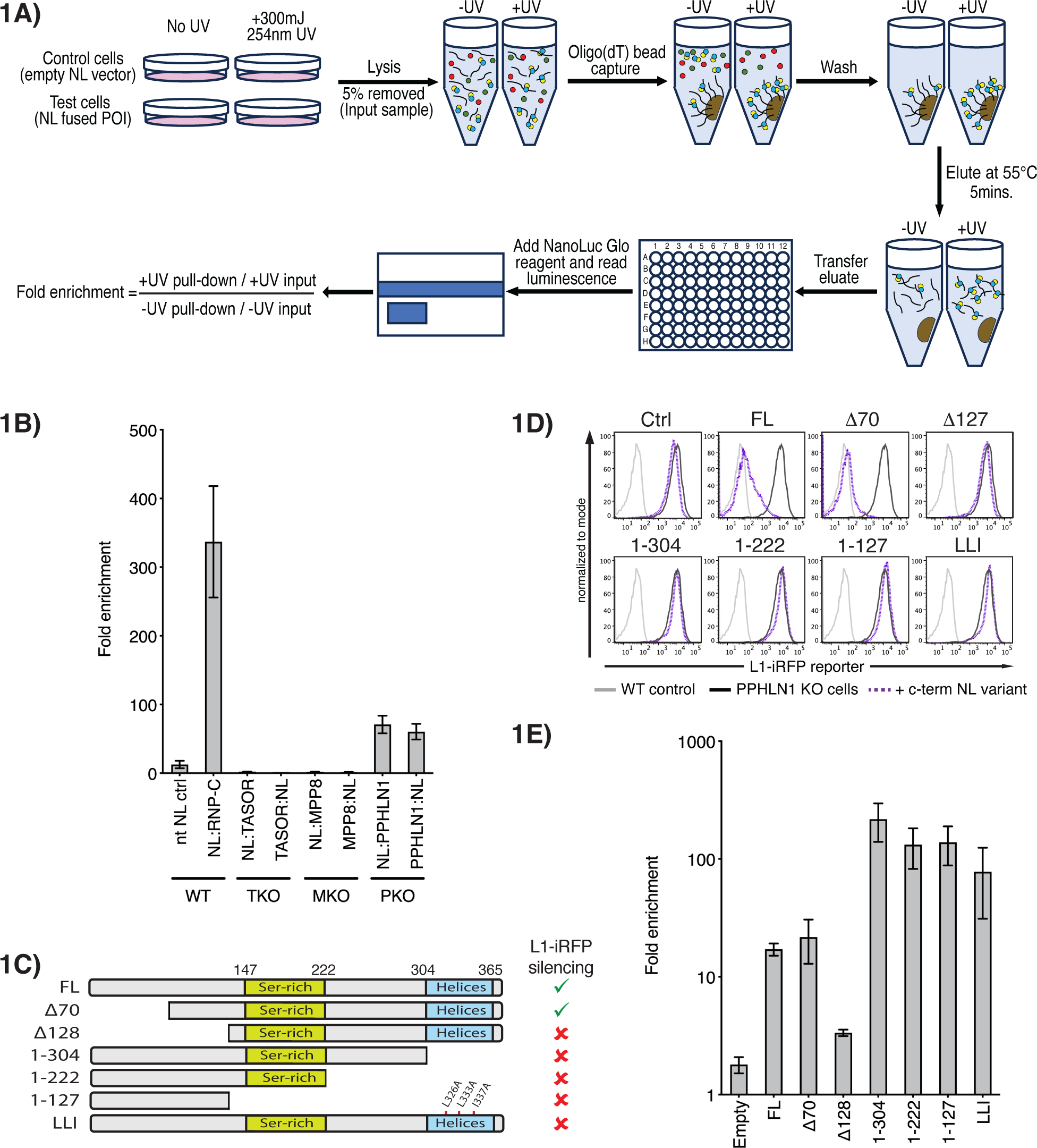
Periphilin is the primary RNA binding protein of the HUSH repressor complex. Experiments performed in HEK293T cells. (**A**) Schematic outlining the steps of the NanoLuc RNA interactome capture assay (NL-RIC). (**B**) NL-RIC assay showing the fold enrichment of UV irradiated samples over non-irradiated samples for both N- and C-terminally NL tagged HUSH proteins (TASOR, MPP8 and Periphilin (PPHLN1). Each assay was performed in the knockout (KO) cells which had been complemented with the relevant HUSH component cDNA (mean of n=2 technical replicates ± SD) (TKO = TASOR KO, MKO = MPP8 KO, PKO = Periphilin KO). Controls were performed in wild-type cells. Empty N-terminal NL vector was included as the negative control and NL:hnRNP-C as the positive control. (**C - E**) Deletion mapping of Periphilin mRNA binding region. (**C**) Schematic of the Periphilin:HA:NL deletion constructs used in (D) and (E). Right sided column summarises the ability of each Periphilin mutant to complement L1-iRFP Periphilin KO reporter cells, with data taken from (D). (**D**) Flow cytometry of L1-iRFP reporter expression in Periphilin KO cells complemented with the indicated constructs. No reporter wildtype control (light grey), L1-iRFP reporter uncomplemented (dark grey) or Periphilin:(NLS)HA:NL deletion complemented (dotted purple). (**E**) NL-RIC RNA binding analysis of the indicated Periphilin mutants in the L1-iRFP Periphilin KO cells, as used in (D), (mean of n=2 technical replicates ± SD).

### Periphilin mRNA binding is directed by its N-terminus and regulated by its interaction with TASOR

The best-characterised Periphilin isoform (variant 2, NM_201515) encodes a 374 amino acid protein **(Fig. 1C)**, with a long, disordered N-terminal domain (NTD) (residues 1 - 127), a serine-rich region (147-222) and a C-terminal helical region (304-368). We recently showed that the long, disordered NTD is essential for silencing but not for HUSH complex assembly (18), while residues within the C-terminal domain (297-374) form the structural interface between Periphilin and TASOR and are essential for both HUSH complex formation and silencing (18). Furthermore, our recent crystal structure of the Periphilin-TASOR interface shows that two Periphilin molecules bind a single TASOR, via Periphilin residues 292–367, with TASOR forming two α−helices that wrap around the outer surfaces of the Periphilin dimer (18). Periphilin mutations L326A, L333A and I337A (LLI) disrupt the Periphilin dimerization interface while the Periphilin L356R mutant targets both Periphilin-TASOR interfaces (18).

To further define the regions of Periphilin required for RNA binding we initially used a mutagenesis-based approach and generated a series of NL C-terminal tagged Periphilin deletion mutants **(Fig.1C)** which were used to complement a Periphilin knockout reporter cell line. By performing both FACS and NL-RIC analysis on the complemented cells, we defined the regions of Periphilin required to silence a HUSH Line-1 (L1-iRFP) reporter **(Fig. 1D)**, whilst also determining the requirements for RNA binding **(Fig. 1E)**. N-terminal truncated Periphilin mutant Δ70 maintained RNA binding and was fully functional **(Fig. 1D/E)**, whilst further deletion of the disordered NTD (residues 1-127) led to a loss of both RNA binding and silencing activity. In contrast, the C-terminal Periphilin deletion mutants (1-304, 1-222, 1-127) which are unable to bind TASOR (18), and the triple point mutant (LLI), which targets the Periphilin interface and prevents Periphilin homodimerization (18), all demonstrated significantly higher binding to RNA than the full length or Δ70 Periphilin constructs **(Fig. 1E)** but were unable to silence the reporter **(Fig. 1D)**. Therefore, Periphilin binds RNA through its N-terminal domain, homodimerizes and associates with TASOR through its C-terminal domain and requires all three functions for HUSH-mediated silencing.

### Artificial recruitment of Periphilin to a HUSH-resistant RNA directs HUSH-mediated repression

An alternative approach to determine the dependence of HUSH-mediated repression on RNA binding was to test whether RNA binding alone was sufficient to trigger HUSH-mediated silencing, or whether other gene-specific factors might be required. We therefore took advantage of the well-characterised bacteriophage interaction between the lambda N peptide (λN, 1-22) and the RNA BoxB stem-loop element (21) (22). A λN-BoxB tethering system was engineered to artificially recruit each individual λN:FLAG-tagged HUSH component (λNF-tagged Periphilin, TASOR or MPP8) to a HUSH-resistant fluorescent reporter into which 12x BoxB stem loops had been inserted **(Fig. 2A)**. Artificial recruitment of λNF:Periphilin to RNA was both necessary and sufficient to induce silencing, an effect not seen with λNF:TASOR, λNF:MPP8 nor λNF:GFP constructs **(Fig. 2B)**. Recruitment of λNF:Periphilin to reporters containing an increasing number of BoxB loop elements allowed titratable silencing of the reporter **(Fig. 2C)**. This ‘engineered’ Periphilin-induced repression remained dependent on all HUSH complex members, as depletion of each individual HUSH component led to de-repression of the fluorescent reporter **(Fig. 2D)** and silencing could not be established in a TASOR KO cell line **(Fig. 2E)**. Importantly, ChIP-PCR showed that recruitment of Periphilin to the reporter was associated with deposition of HUSH-dependent H3K9me3 over the repressed reporter **(Fig. 2F)**. Silencing by λNF:Periphilin in the BoxB tethering system was also independent of: (i) the cell line used (**Fig. S2B**) (ii) the promoter used to drive the reporter (**Fig. S2C**), (iii) the delivery system, as PiggyBac transposon vectors work as well as lentiviruses (**Fig. S2D**), (iv) the tethering system used, as the MS2:MS2 coat protein (MCP) system is as effective as the λN-BoxB (**Fig.S2E**), or (v) the reporter protein used (**Fig. S2C and Fig. S2F**). These experiments provide further evidence that RNA-binding by Periphilin is required for HUSH-dependent repression.

**Fig. 2.**
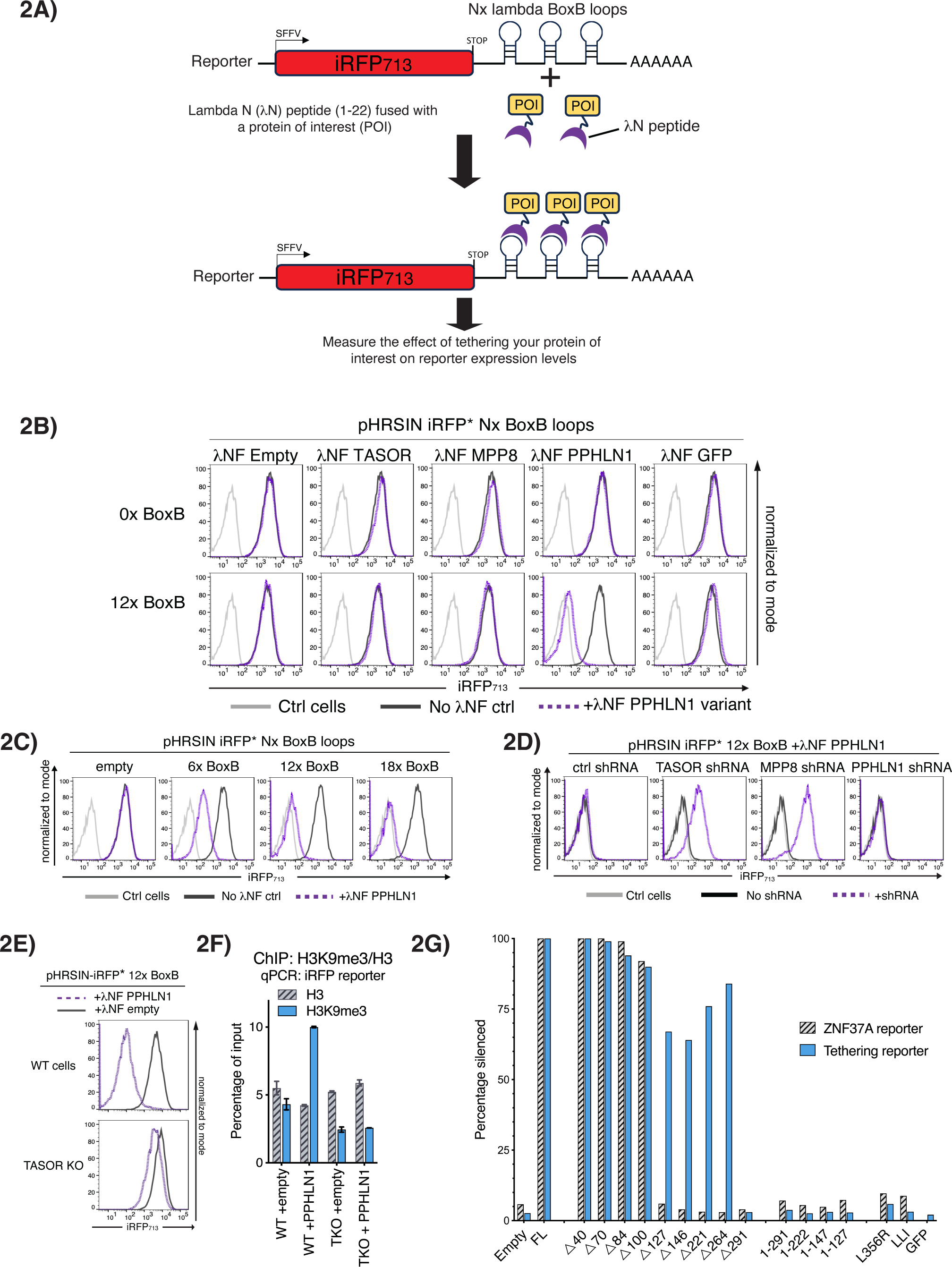
Recruitment of Periphilin to RNA bypasses the RNA binding requirements of Periphilin. Experiments performed in Jurkat cells. (**A**) Schematic of the Lambda N (λN)-BoxB stem loop tethering system. The λN(1-22):FLAG tag fused protein of interest (POI) is recruited to BoxB stem loops inserted downstream of a HUSH insensitive iRFP reporter, and used to monitor reporter expression. (**B**) Recruitment of λNF_tagged HUSH complex components to 0x and 12x BoxB reporter systems. Wildtype cells (light grey), reporter cells with no λNF construct (dark grey) or with λNF fused POI (dotted purple). (**C**) Tethering assay showing the effect of increasing the number of BoxB hairpin loops, Wildtype cells (light grey), λNF empty (dark grey) or with λNF fused POI (dotted purple). (**D**) shRNA knockdown of individual HUSH components in a iRFP* 12x BoxB + λNF:Periphilin silenced reporter Jurkat cell line shows HUSH dependency of the Periphilin tethering (control shRNA is non-targeting, the λNF:Periphilin construct is resistant to the Periphilin targeting shRNA). Wildtype cells (light grey), no shRNA (dark grey) and plus shRNAs (dotted purple). (**E**) Transduction of λNF:Periphilin into wildtype vs TASOR knockout BoxB reporter cells, shows the requirement for TASOR to establish silencing. λNF empty (dark grey) or with λNF:Periphilin (dotted purple). (**F**) H3K9me3 ChIP-qPCR analysis of samples from (E) showing H3K9me3 deposition on reporter in λNF:Periphilin expressing but not TASOR KO cells. (**G**) Graphical representation of the percentage of cells showing significant silencing in Jurkat Periphilin knockout cells harbouring dual ZNF37A and iRFP* 12x BoxB tethering reporters, following complementation with λNF(NLS):Periphilin deletion and mutant constructs (Fig. S2H). ZNF37A reporter (grey diagonal stripe) and iRFP12x BoxB tethering reporter (solid blue).

### Artificial recruitment of Periphilin to RNA bypasses the RNA binding requirements of Periphilin

The previous experiments showed that artificial recruitment of Periphilin to an RNA transcript can silence an otherwise HUSH-resistant reporter. We next wanted to determine whether the BoxB-mediated recruitment of λNF:Periphilin could bypass the RNA-binding requirements of Periphilin. Using a series of N-terminal Periphilin deletion mutants we confirmed that, in the absence of tethering, deletion of the first 127 amino acids of Periphilin results in a Periphilin mutant unable to silence a HUSH-sensitive reporter located within the ZNF37A gene of a Periphilin knockout cell line **(Fig. 2G and Fig. S2G upper panel)**. In contrast, following the BoxB mediated tethering of Periphilin to a specific transcript, the first 264 N-terminal amino acids of Periphilin could be removed, leaving only a minimal 110 amino acid λNF:Periphilin Δ264 mutant (amino acid residues 265-374) that was still able to maintain ∼85% HUSH repression **(Fig. 2G and Fig. S2G lower panel)**. This minimal λNF:Periphilin Δ264 domain (a.a. 265-374) is devoid of RNA-binding capacity but includes the residues critical for Periphilin homodimerization and binding to TASOR (18).

We confirmed that the Periphilin point mutant that disrupted the Periphilin-TASOR interface (L356R), as well as the Periphilin dimerisation mutants (LLI) were unable to repress the ZNF37A HaloTag reporter in Periphilin KO cells **(Fig. 2G and Fig. S2G upper panel)**. Furthermore, BoxB-mediated recruitment of these same λNF:Periphilin mutants also failed to silence the HUSH-resistant iRFP BoxB fluorescent reporter **(Fig. 2G and Fig. S2G lower panel)**. Therefore, despite bypassing the requirement for RNA binding, artificially tethered Periphilin must still homodimerize and bind TASOR to enable HUSH-mediated silencing.

### The HUSH complex can mediate silencing via nascent RNA

These experiments show that HUSH-mediated repression is RNA-dependent but do not distinguish binding to nascent as opposed to processed RNA transcripts. By inserting BoxB elements within an intron we were able to test whether recruitment of HUSH to a spliced intron, and therefore to nascent (pre-spliced) RNA **(Fig. 3A)**, was sufficient to establish HUSH-dependent repression. In the presence of λNF:Periphilin, insertion of 12x BoxB elements into the intron of a HUSH-resistant iRPFi reporter induced HUSH-dependent silencing of the reporter **(Fig. 3B)**, as confirmed by the HUSH-dependent deposition of H3K9me3 across the reporter construct **(Fig. 3C)**. Genomic and RT-PCR analysis using primers that flanked the intron, confirmed appropriate splicing of the construct with no splicing defects caused by either HUSH recruitment or insertion of the BoxB elements **(Fig. S3A)**. Furthermore, using an iRFP reporter **(Fig. 3D)** and positioning the intron either (i) within the 5’UTR (ii) internal to iRFP cDNA, or (iii) within the 3’UTR of the transcript, resulted in efficient λNF:Periphilin-mediated silencing by all three reporter constructs, though silencing was most efficient when tethered within the 3’UTR of the transcript **(Fig. 3D)**. Periphilin binding to nascent RNA transcripts can therefore induce HUSH-dependent silencing.

**Fig. 3.**
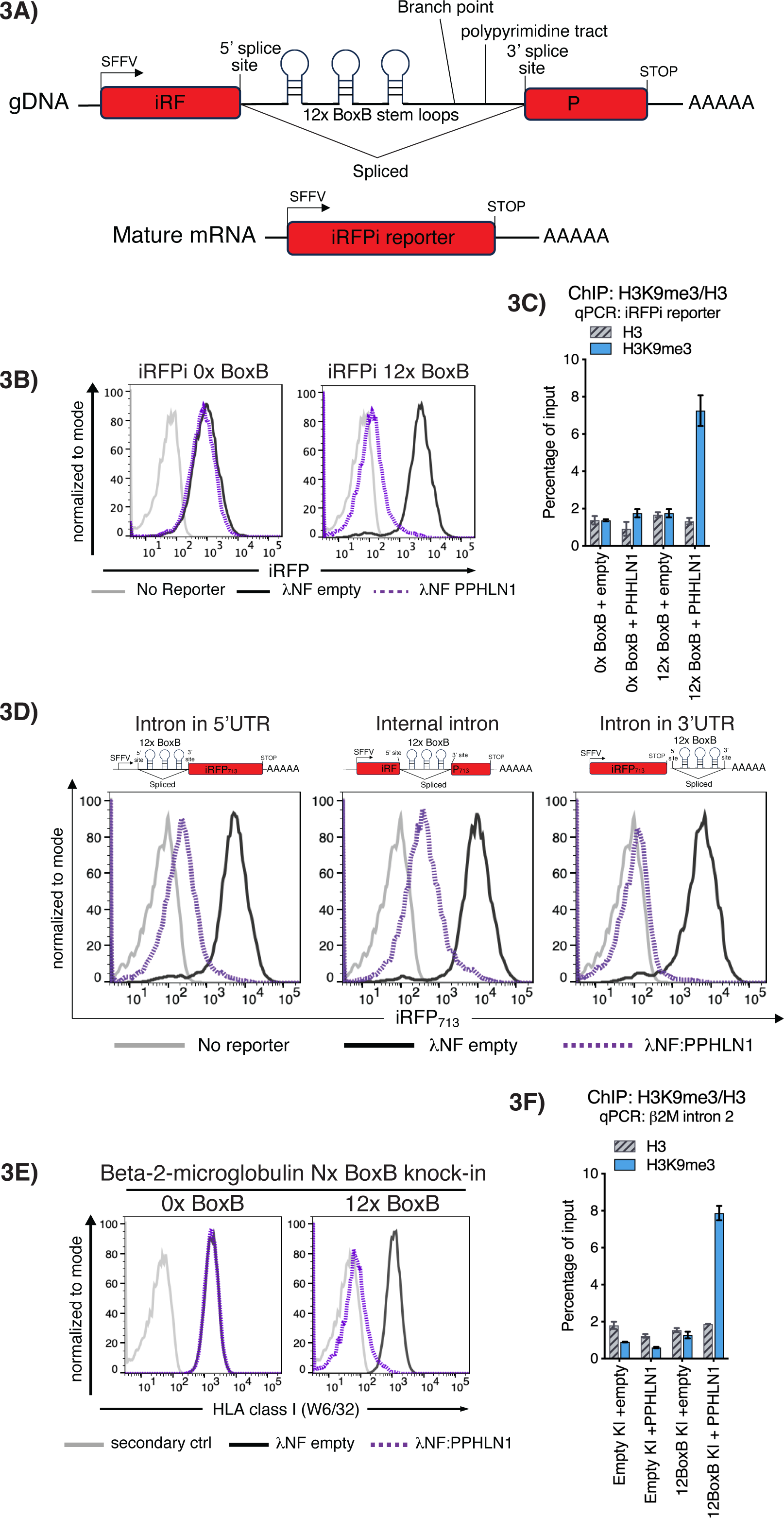
The HUSH complex can mediate silencing via nascent RNA. Experiments performed in Jurkat cells. (**A**) Schematic of λN-BoxB tethering system inserted into an intron. An U2 dependent, 12x BoxB stem loop containing intron was introduced into a iRFP reporter construct (iRFPi) (**B - D**) Recruitment of λNF:Periphilin to the iRFPi reporter system is sufficient for reporter silencing. (**B**) Flow cytometry of λNF empty and λNF:Periphilin transduced iRFPi containing 0x BoxB and 12x BoxB reporter cells. (**C**) H3K9me3 ChIP-qPCR analysis of samples in (B) (**D**) Flow cytometry of iRFPi 12x BoxB reporter constructs showing intron-tethered λN-BoxB silencing is independent of the intron location. Intron locations, 5’UTR (**left panel**), iRFP internal (**centre panel**) and 3’UTR (**right panel**). Wildtype cells (light grey), reporter cells with λNF empty (dark grey) or with λNF:Periphilin (dotted purple). (**E**) Flow cytometry of λNF empty and λNF:Periphilin transduced Jurkat cells, expressing BoxB stem loops inserted into the 3’ end of the beta-2-microglobulin (β2M) locus. Secondary only stained wildtype cells (light grey), λNF empty (dark grey) or with λNF:Periphilin (dotted purple). (**F**) H3K9me3 ChIP-qPCR analysis of samples in (E) showing H3K9me3 deposition on β2M intron 2, in λNF:Periphilin vs. λNF empty transduced cells.

### Silencing of the beta-2-microglobulin gene by artificial recruitment of Periphilin to its RNA

Our previous experiments demonstrate how tethering Periphilin to a reporter transcript can silence its expression. To determine whether BoxB-λNF:Periphilin can be used for the HUSH dependent silencing of an endogenous gene, we used CRISPR-Cas9 and donor plasmids carrying either zero or twelve BoxB loop elements to target the 3’ UTR of the endogenous beta-2-microglobulin gene (β2M) in the Jurkat cell line. **(Fig. S3C and Fig. S3D)**. The introduction of λNF:Periphilin to the 12x BoxB knock-in cells effectively reduced cell surface MHC-I expression, due to β2M silencing, and was not seen in the 0x BoxB knock-in cell line **(Fig. 3E)**. ChIP-PCR analysis showed that this silencing also resulted in HUSH-dependent H3K9me3 deposition over the repressed β2M gene **(Fig. 3F)**.

### Periphilin RNA binding specificity is independent of its C-terminus or other HUSH components

To characterise further the endogenous targets of Periphilin binding we performed UV-crosslinked RNA immunoprecipitation sequencing (UV-RIPseq) in Periphilin and SETDB1 dual knockout HEK293T cells, expressing a panel of HA-tagged Periphilin constructs. The dual KO cells were used to prevent regulation of the functional Periphilin mutants by HUSH, which therefore allowed for more comparable Periphilin mutant expression levels **(Fig. S4A)**. UV-crosslinking ensures maintenance of the protein:RNA interaction during subsequent downstream processing. Following HA-bead immunoprecipitation, the RNA bound to the immunoprecipitated protein was extracted for subsequent analysis. We performed UV-RIPseq on empty vector, full length Periphilin, the Periphilin truncation mutants 1-127, 1-147, and 1-222, and the Periphilin point mutants L356R and LLI.

Full length Periphilin protein bound the previously described HUSH silencing targets, e.g. L1s and ZNF genes (**Fig. 4A**) (7). To determine if the Periphilin mutants have the same RNA binding specificity, we used the R package DiffBind to identify sites of differential binding between the ∼31,000 peaks identified in the full length and mutant Periphilin constructs. This showed that Periphilin mutants 1-147, 1-222, L356R and LLI all phenocopy the wildtype Periphilin profile, whereas mutant 1-127 binds approximately 10% less HUSH-associated sites (P value 8.28 x 10^-209^) (**Figs. 4B - D**). This effect is most likely due to the reduced expression of Periphilin 1-127 compared with the other constructs **(Fig. S4A)**, though we cannot exclude a true biological effect. The RIP-seq data was also examined to determine which genes, whose transcripts were bound by Periphilin, were rich in transposable element (TE) **(Fig. 4E)**. Compared to genes in reference genome (GRCh38), Periphilin-bound genes were markedly enriched in TEs, irrespective of which mutant was tested. These data would suggest that the specificity of Periphilin RNA binding is independent of its C-terminus and therefore TASOR binding.

**Fig. 4.**
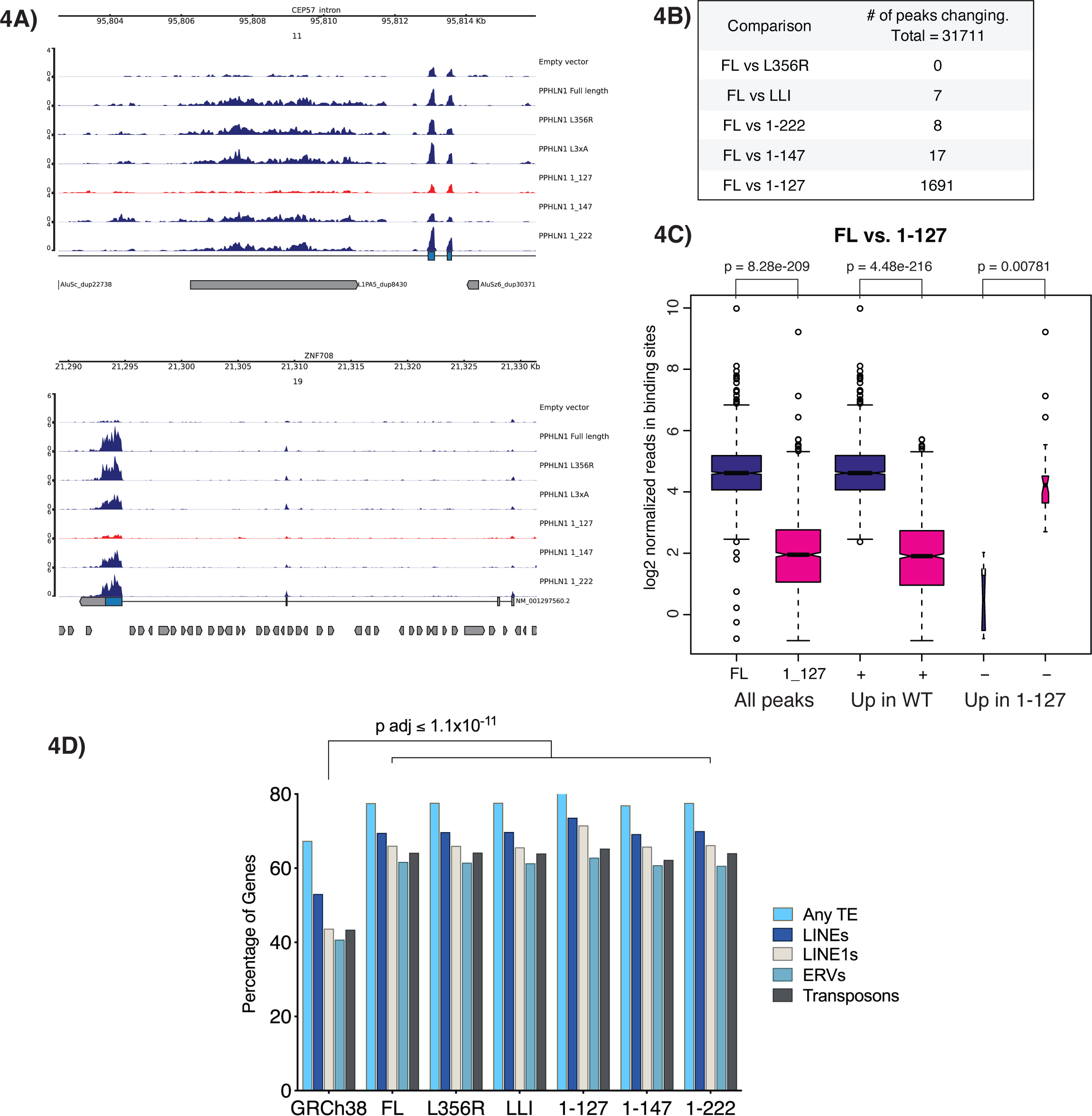
Periphilin RNA binding specificity is independent of its C-terminus or the other HUSH components. RIP-seq analysis performed in Periphilin and SETDB1 dual knockout HEK293T cells (**A**) Genome browser tracks showing Periphilin enrichment over the CEP57 intronic Line-1 and across the ZNF708 gene coding regions. (**B**) Analysis of the number of differential RNA-binding peaks for each of the indicated Periphilin variants as compared to the 31,711 peaks identified for full length (FL) Periphilin. (**C**) Box plot showing normalised binding site reads for FL Periphilin vs Periphilin truncation mutant (1-127) pulldowns. (**D**) Transposable element (TE) mapping within transcripts bound by Periphilin shows enrichment for transposable elements within Periphilin bound genes. Adjusted p-values determined by Tukey multiple comparisons of means, showing a significant difference between the TE content in all Periphilin variants compared to all genes in the genome. See supplementary data for a full statistical comparison.

## Discussion

The identification of HUSH as an epigenetic repressor complex able to identify and silence any newly integrated retroelement raises important questions as to how HUSH recognises such a diverse array of unrelated DNA sequences. Our finding that HUSH uses the presence and absence of cellular introns to differentiate ‘self-DNA’ (intron-containing) from ‘non-self DNA’ (retroelement-derived) provided important clues as to how HUSH distinguishes its targets, but understanding how this might work mechanistically remains a challenge (8).

An essential role for RNA in HUSH-dependent silencing arose from the observation that active transcription is required for HUSH recognition. Together with the observed similarities between HUSH and the RITS silencing complex, the finding that silencing of identical DNA sequences is directional, relative to RNA Pol II transcription, together with the association of TASOR with components of the RNA processing machinery (9,12,19) all pointed to a central role for RNA in HUSH-dependent silencing. While none of the three core HUSH components contains an identifiable RNA-binding domain, each HUSH component is reported to have some *in vivo* RNA-binding capability (9,12,15,17-19), though how much of this binding could be ascribed to an individual protein as opposed to the whole HUSH complex has been unclear. Equally important to establish was whether RNA binding is essential for HUSH function, given the proximity of the HUSH complex to the chromatin environment and sites of transcription.

In this study we took an unbiased approach and found that Periphilin was the only HUSH component to demonstrate significant RNA binding, that its RNA-binding region lies within the first 147 amino acids of Periphilin and this region is essential for HUSH function. By artificially recruiting individual HUSH components to RNA reporters we showed that not only is Periphilin the only HUSH component able to repress an otherwise HUSH-insensitive reporter, but that Periphilin binding to nascent unprocessed RNAs can initiate HUSH-dependent repression. By tethering Periphilin to a specific transcript we were able to by-pass the need for Periphilin’s RNA binding domain completely. Under these conditions the only region of Periphilin required for HUSH activity was its 110 amino acid C-terminal domain involved in homodimerization and binding to TASOR.

These data, together with our published genome-wide Periphilin RIP-seq analysis showing that transcripts bound by Periphilin are known HUSH targets (full-length L1s, KRAB-ZNFs and HUSH-sensitive loci) (7), strongly suggest that Periphilin binds nascent RNAs, and these RNAs specify the loci for HUSH repression, most likely through directing and stabilising HUSH at their target loci. Our current model of HUSH therefore puts Periphilin binding to RNA as an early event in the initiation of HUSH-dependent silencing. HUSH may track RNA Pol II and upon triggering of HUSH, as will occur if RNA Pol II is transcribing through an unusually long, A-rich exon, Periphilin will bind the nascent transcript and recruit the other HUSH complex components to the locus to be silenced.

While some features of HUSH are reminiscent of other well-characterised RNA-dependent silencing complexes, such as the requirement for transcription to initiate and maintain silencing, and the ability to bind nascent RNA(13,23-25), there are also unique features of HUSH-dependent silencing which need to be explained. In particular, HUSH has no clear requirement for processing small RNAs, for Argonaute-like binding proteins, nor does it have a mechanism for amplifying the RNA signal. While RITS uses its RNA-dependent RNA polymerase (RdRP) to amplify and reinforce the RNA silencing signal (26,27) and piRNA silencing systems use a ping-pong amplification cycle (28,29), no such mechanism has been identified in silencing by HUSH, nor do vertebrates encode the RdRP enzyme. How HUSH compensates for this deficit remains unclear. Even when the RNA binding requirements of Periphilin are bypassed, by artificially recruiting Periphilin to a specific transcript, the minimal 110 amino acid Periphilin fragment must still retain its ability to homodimerize, bind TASOR and assemble into the HUSH complex. This may reflect a requirement for Periphilin to oligomerize and form higher order molecular condensates (18), potentially as a means of reinforcing the RNA signal.

An unexpected finding was that all the Periphilin C-terminal deletion mutants as well as point mutants which prevented Periphilin from binding to TASOR showed significantly higher RNA binding compared to full length Periphilin and could not be explained by differences in Periphilin expression levels. Periphilin binding to RNA therefore appears to be decreased when dimeric Periphilin is incorporated within the HUSH complex. While we don’t fully understand this observation, it may reflect an innate ability of free Periphilin to bind RNA, with this function moderated following Periphilin’s assembly within the complex. This suggests a model in which Periphilin binding to RNA is initiated by an upstream event, for example a change in transcriptional activity which is sensed by HUSH and triggers Periphilin to bind RNA. These details require further investigation but are supported by our RIP-seq analyses showing that Periphilin binds similar RNA species whether incorporated within the HUSH complex or not bound to TASOR. Future studies will need to determine whether Periphilin has a clear RNA binding preference/sequence specific motif, or whether binding is promiscuous and a response to a dynamic change in HUSH behaviour. These studies would be facilitated by *in vitro* studies of RNA binding by Periphilin, but the difficulties in expressing full-length HUSH complex proteins *in vitro*, has made the direct assessment of RNA binding capacity challenging.

The possibility of regulating endogenous genes by recruiting Periphilin to a specific transcript, as shown with the beta-2-microglobulin locus, demonstrates a potential use for this technology in regulating endogenous genes. This could be used to repress essential genes whose deletion is otherwise lethal and has the advantage of not requiring manipulation of the genome itself. As HUSH is transcription dependent, gene expression is unlikely to be completely repressed and the degree of repression can be titrated by adjusting the number of BoxB elements in the transcript. The inclusion of an inducible system for expression of the lambdaN:Periphilin component could further enhance the flexibility of regulation, and using a Periphilin NTD deletion mutant would avoid perturbation of normal Periphilin/HUSH function.

In conclusion our results emphasise an essential requirement for Periphilin in binding RNA and initiating the repressor functions of the HUSH complex, and implicate the RNA-binding features of Periphilin as a key determinant of HUSH specificity in defining its substrates.

## Acknowledgements

We thank all members of the Lehner lab and Professor Yorgo Modis (University of Cambridge Department of Medicine) for critical discussions. We thank Professor Anna Petrunkina-Harrison and the CITIID Flow Cytometry Core Facility team and the NIHR Cambridge Biomedical Research Centre (BRC). For the purpose of open access, the University of Cambridge has applied a CC BY public copyright licence to any Author Accepted Manuscript version arising. This work was supported by a Wellcome Trust Discovery Award (227418/Z/23/Z) to PJL.

